# Knowledge production and translation of science between Academia and Industry: assessing the impact of R&D in India

**DOI:** 10.1101/2021.05.08.443271

**Authors:** Vishal Rao US, Naga Durga Ram Jenu, Gururaj Arakeri, Ravi C Nayar, Jitendra Kumar, Rui Amaral Mendes

## Abstract

India is at the 3^rd^ position worldwide in terms of publication of scientific literature. However, in terms of productivity, it has been consistently failing to transform the research knowledge into industrial output. This study compares India with the leading countries to understand its lacuna in terms of R&D policy and outputs. Although scientific publications are regarded as the output of basic research, patent applications serve as a better indicator of the applied research. This paper assesses the important determinants for patent filings of a nation. It also focuses on the role of academia and industry collaboration in R&D and the productivity of a nation. We found that the higher the GERD (total Gross Domestic Expenditure of R&D) and the R&D personnel in a nation, the higher the patent filings of the nation. Moreover, we show that academia-industrial collaboration plays a key role in transforming basic research into real-world applications, as we illustrate the government’s role in making necessary policies to make the collaboration successful. This paper highlights the significance of investing in R&D to improve the productivity of a nation, as also the need to design policies to strengthen the applied research environment by fostering solution-centric collaborations between academia and industry.

## Introduction

Knowledge production is changing, as are societies’ long-term and broader expectations from that knowledge. This milieu places academia on a pivotal pedestal on the cusp of teaching, research, and “third mission”, the latter relating to activities that are expected to propel and enhance the social, economic and cultural development, both by transferring knowledge and technologies to industry and the society at large (1).

The translation of rapidly evolving scientific knowledge into healthcare practice, organization, and policy is a daunting challenge worldwide, often related to how different stakeholders perceive their roles in the transference of knowledge (2).

Looking at the changes in the third mission profiles of universities in the United Kingdom and how these evolved in terms of knowledge exchange activities and partnerships, Sanchez-Barrioluengo found a very diversified set of profiles of third mission activities across different types of universities, ranging from research-oriented activities typically in partnership with large firms and non-commercial organisations to relatively less research-intensive universities relying on consultancy and formation of spin-offs (2). Since, research is crucial to the technical advancement of industries and innovations, a developing nation like India should encourage more strategies and policies which foster innovation capabilities.

Although, India is the 5^th^ largest economy in terms of GNI (Gross Net Income) after the USA, China, Japan and Germany respectively (Figure 1). It is classified under the lower-middle-income category in terms of GNI per capita (US$ current). It is also the only BRICS country to fall under this category (3).

**Figure 1.**
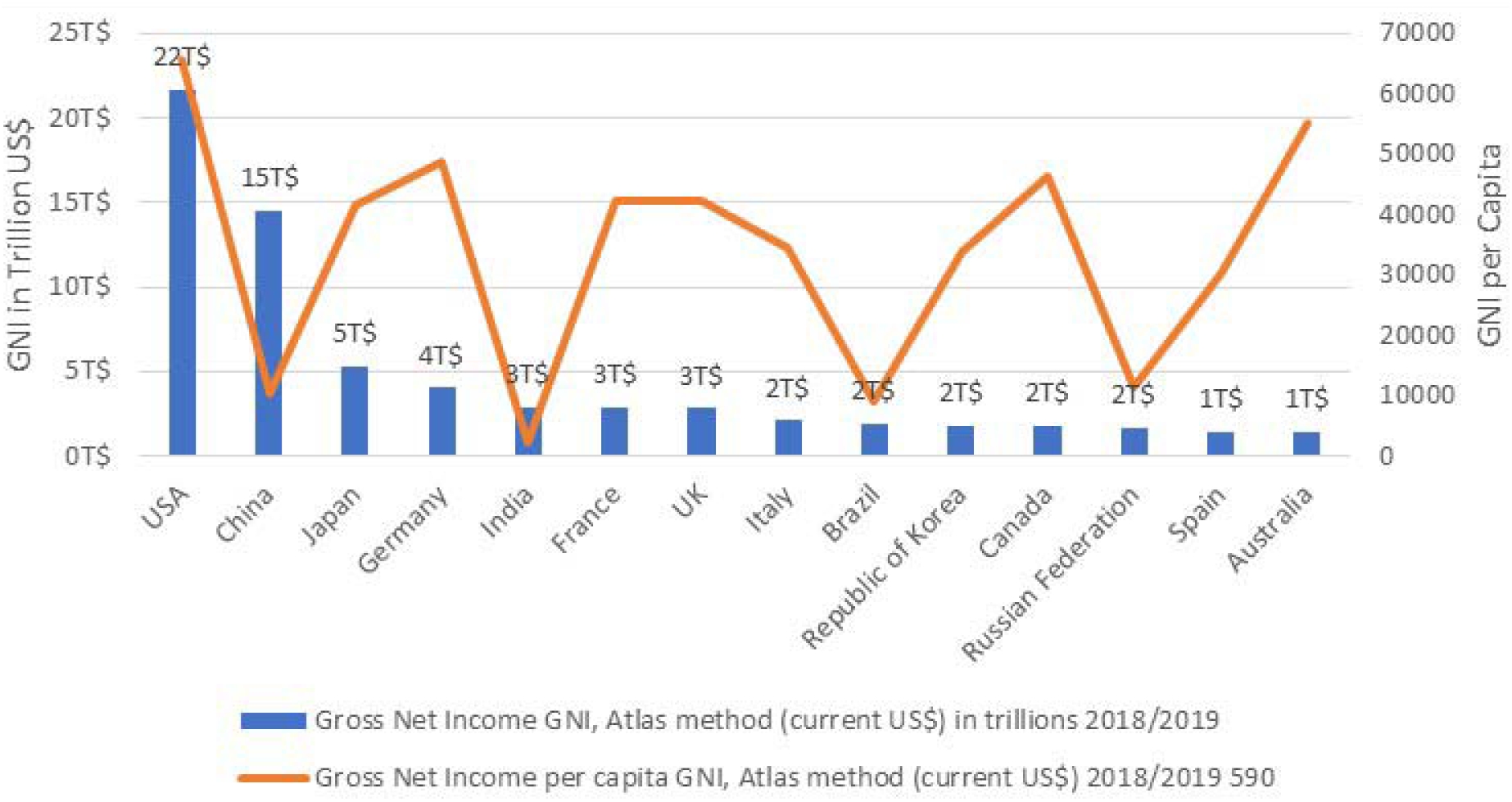
Comparison of Gross Net Income (GNI) & Gross Net Income per Capita of top countries as per 2019 or recently available data.

However, India has a bold aspiration to become an upper-middle-income country by achieving a US$10 tr GDP by 2034. In order to achieve this, India needs to grow its GDP at a compound annual growth rate (CAGR) of 9% over the next 20 years (4) but the steep slopes in the graph line (Figures 1) show that the GNI per capita for the BRICS countries is very low compared to that of other leading or developed countries. This is due to their predominant dependency on exporting less value-added products. High-value addition comes with invention and innovation, which in turn comes with highly skilled personnel and a research fostering environment. Although India has the second-highest working population in the world after China (5), it faces an employment deficit in terms of skills (6). Contrary to popular belief, only 18% of global goods trade is driven by labour cost, Thus, making low-skill labour a less important factor of production (7). The possible breeding grounds for upskilling and innovation in India are higher education institutions but only 1% of them are involved in (R&D) Research and Development. (8). In essence, it may that India need to focus more on developing its human resource potential and creating a fostering environment for research and innovation.

The countries which are leading in terms of innovation have been investing their resources in R&D and also laid policies accordingly to create a fostering environment to produce high value-added products. In terms of research publications and patent filings, the neighbouring country China has been performing outstandingly not only among the BRICS countries but also among the developed nations; a similar trend has been observed in South Korea as well (9).

Interestingly, both China and South Korea were once agrarian countries where the economy was purely based on agriculture exports but both transformed into innovation leaders of the world (9). South Korea, in particular, has moved from Asia’s poorest to one of the wealthiest countries (10). These two countries have so much in common and India can learn from their experience and change its strategy and policies, to become one of the innovation leaders of the world (9). Otherwise, it may likely remain a lower-middle-income country for another generation.

If we analyse the history of China & the Republic of Korea, both have gone through two major economic transitions, one transition is from agrarian exporter to industrial manufacturing around the 1980s and the second transition from industrial manufacturing to innovation centres. In both the transitions, respective governments played a significant role by bringing policy changes to enhance the collaboration between academia and industry during the 1990s for the overall benefit of the economy (8). Worldwide, the most Triple-Helix model (Figure 2) became successful in which major stakeholders are academia, industry and government. Here, academia brings the knowledge to address the research problems of the industry. Industry brings skills, technology and funds required by the academia to address the research and infrastructure problems of the academia and the government brings support in terms of funds and facilitating policies between the academia and industry and foster the innovation ecosystem and improve its economy (11).

**Figure 2.**
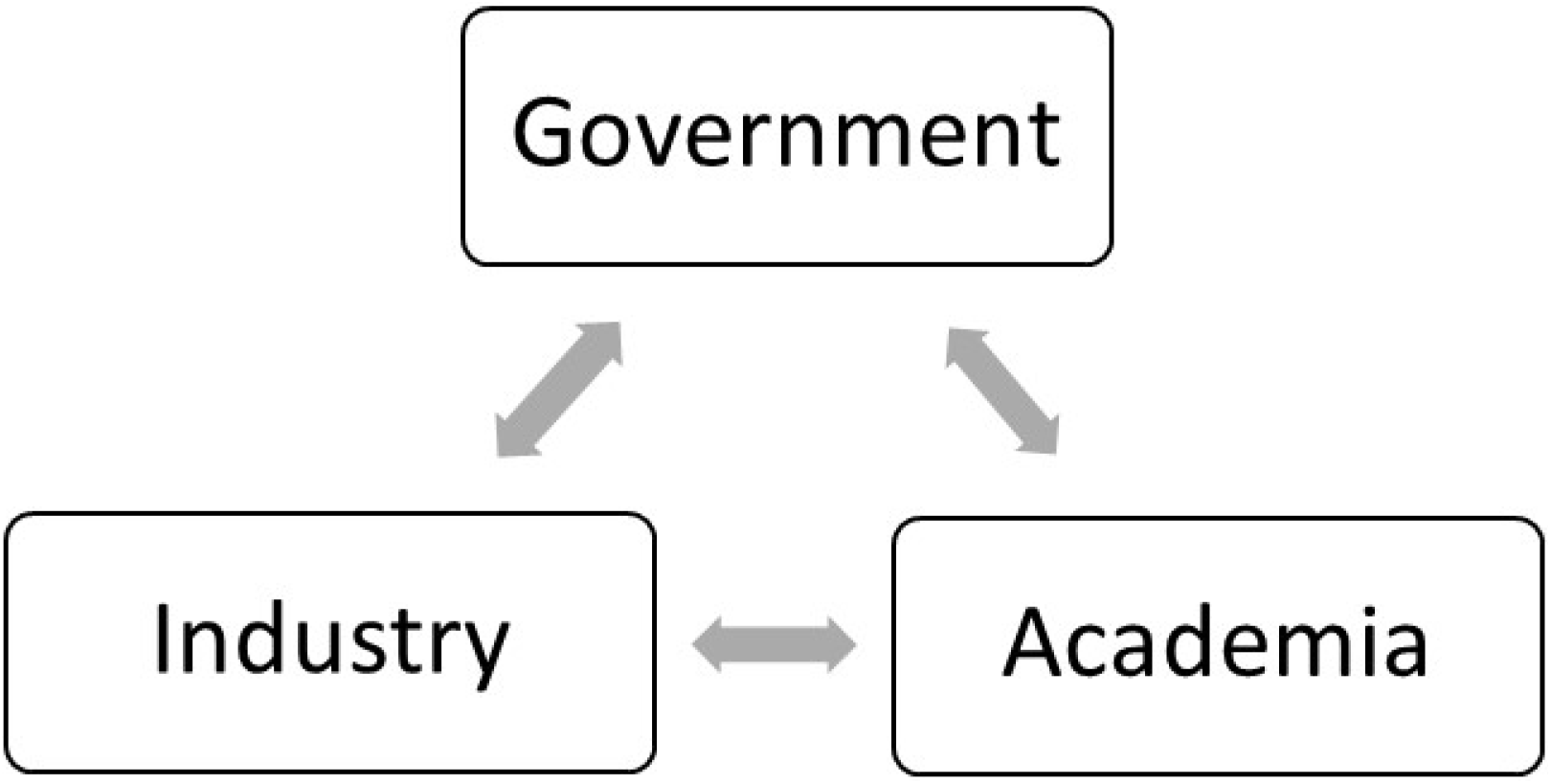
Triple Helix Model

Although India has been trying to improve the industry output by the Make-In-India initiatives, most of the policies are not fulfilling (12). A huge skill gap in the working population making it hard to compete against China, even with other south Asian countries like Thailand, Indonesia etc. Many manufacturers in India had never given a thought of invention before. If the local manufacturers can get research capabilities from the higher education institutes and invest in R&D, they surely improve the quality and produce patented products, but Indian education institutes often have a low level of research output. In contrast, China has made a strenuous effort to revamp its higher education sector since the late 1990s as part of its five-year plans (9). It increasingly focussed on research and gradually outstripped the US in terms of scientific research publications and patent filings (Figure 5).

Scientific publications and patents are globally accepted indirect efficiency output indicators of innovation (13). In this study, we critically analyse the world data and try to identify whether spending on R&D and increasing R&D personnel will improve the industry and academia performance in terms of scientific publications and patents and whether India should change its strategy and policy towards R&D capabilities to improve its economy stance in the world.

## Materials and Methods

More than 100 measurable indicators were checked for face validity to measure the construct is supposed to measure. Out of them, few indicators were selected to achieve the objectives of this research. A quantitative correlational approach was adopted with a total of 163 countries available data and the top leading countries have been shortlisted by selecting the countries with top patent filings on average of the last five years available data.

GNI dataset was extracted from the World Bank (Last updated on 02, 2021). The reason why GNI has been chosen over GDP is GNI measures the total income received by the country from its residents and businesses regardless of whether they are located in the country or abroad. Countries like India have the largest diaspora, where they have large foreign receivables or outlays. Therefore, GNI would be a better choice than GDP.

R&D personnel (Researchers, Technicians & supporting staff) count was extracted from the UIS (UNESCO Institution of Statistics) on March 1st, 2021. In this study, the recently available data was selected for each country. To most of the countries, 2018 data is available. For those countries where data for 2018 is not available, previous year data was selected. In UIS, there are two variants of DATA is available about R&D personnel, one is FTE (Full Time Equivalent) and HC (HeadCount). In this study Headcount variant was chosen because FTE may not show the actual number of personnel that exists in a nation, whereas the HeadCount would denote the actual number of R&D personnel.

GERD (Gross Domestic Expenditure on R&D) data was also extracted from the UIS on March 1st, 2021. GERD total is available in three variants i.e., local currency, PPP$ (current) & PPP$ which is pegged to constant prices of 2005. Since local currencies make comparison difficult, PPP$ (current) is used in the study in which the exchange rate equalizes the purchasing power of different currencies which makes it possible for international comparison of current economies.

GERD by source of funds (total intramural) Dataset was extracted on 22 Mar 2021 from UIS as well and took the average of last five years available data. It covers different sectors of the economy (business enterprise, government, higher education, private non-profit organizations) or from “the rest of the world”.

GERD by sector of performance and field of research data was extracted from the OECD database on March 24, 2021. This dataset comprises research and development (R&D) expenditure by sector of performance (business enterprise, government, higher education, private non-profit, and total intramural) and by field of research (natural sciences, engineering and technology, medical and health sciences, agricultural and veterinary sciences, social sciences, and humanities and the arts).

Sector-wise business enterprises expenditure on R&D (BERD) has been extracted from the European Commission which was updated on 1st Jan 2020 by the time it is being extracted. The original data was represented in Euros currency which is converted to USD dollars currency using the current rate i.e., on Mar 30th 2021. At the rate of 1 Euro = 1.17 USD dollar.

Patents data was extracted on 03 Mar 2021 from WIPO (World Intellectual Property Organization) statistics database (Last updated: January 2021). The total number of patents covers solely the filings in each country and doesn’t specify grants of the patents. It is calculated by the applicant’s residence or nationality. This data comprises both residents and abroad patent filings by the residents of each country and excludes the non-resident applications from their respective countries. To avoid the fluctuations, the average of the last five years data is used for analysis i.e., from 2015-2019 years.

Publications data was extracted on 25 Feb 2021 from National Center for Science and Engineering Statistics, National Science Foundation; Science-Metrix; Elsevier, Scopus abstract and citation database, accessed June 2019. However, Articles may have co-authors from different countries, which force us to opt for the “fractional count” over the “whole count” to reflect the actual contribution and to minimise the fluctuations, the average of the last five years available data is used for analysis i.e., 2014-2018 years.

## Results

UNESCO (United Nations Educational, Scientific and Cultural Organization) along with The OECD (Organisation for Economic Cooperation and Development) and EUROSTAT developed an indicator GERD (Gross Domestic Expenditure on R&D) which is expressed as a percentage of Gross Domestic Product (GDP) for international comparison (Indicators of Sustainable Development: Guidelines and Methodologies, 2007). Israel had spent the highest with 4.9% followed by the Republic of Korea with 4.8%. Whereas India stood at 38th place with 0.64% of GDP.

Though many countries use this ratio frequently in their statistics as an indication of the level of financial resources devoted to R&D, it doesn’t necessarily indicate the outcomes of it especially, when comparing developing countries with huge population & economic disparities (Frascati Manual 2002, 2002). Though Israel spends the highest GERD as a percentage of GDP, still its expenditure is 32 times less than that of the USA (United States of America) (14). Hence, Instead of GERD as a percentage of GDP, the total GERD itself would make sense to measure and compare the innovative outcomes of a country.

A regression analysis between the Total GERD and Patent filings of a country has shown that GERD has a significant impact on the patent filings in a country but itself may not be the sole influencing factor. In the case of the USA & China, both countries spent outstandingly highest than other countries but China had filed two times more patents than the USA in 2018. Germany had spent more than the Republic of Korea but fell behind it when it comes to Publications and patent filings. According to the 2018 dataset of UIS, it was found that the other important influencing factor is total R&D personnel in a country. Total R&D personnel has shown a significant impact on patent filings in a country.

The significance became even stronger when total R&D personnel plotted along with the total expenditure of a nation about the total number of patent filings of country. In a multiple regression analysis (Table 1), a strong linear relationship has shown that the two factors expenditure and R&D personnel together have a statistically significant impact on the patent filings in a country. In other words, the higher the innovation inputs like GERD and R&D personnel higher the innovation outputs like patent filings of a country (Figure 3).

**Table 1.**
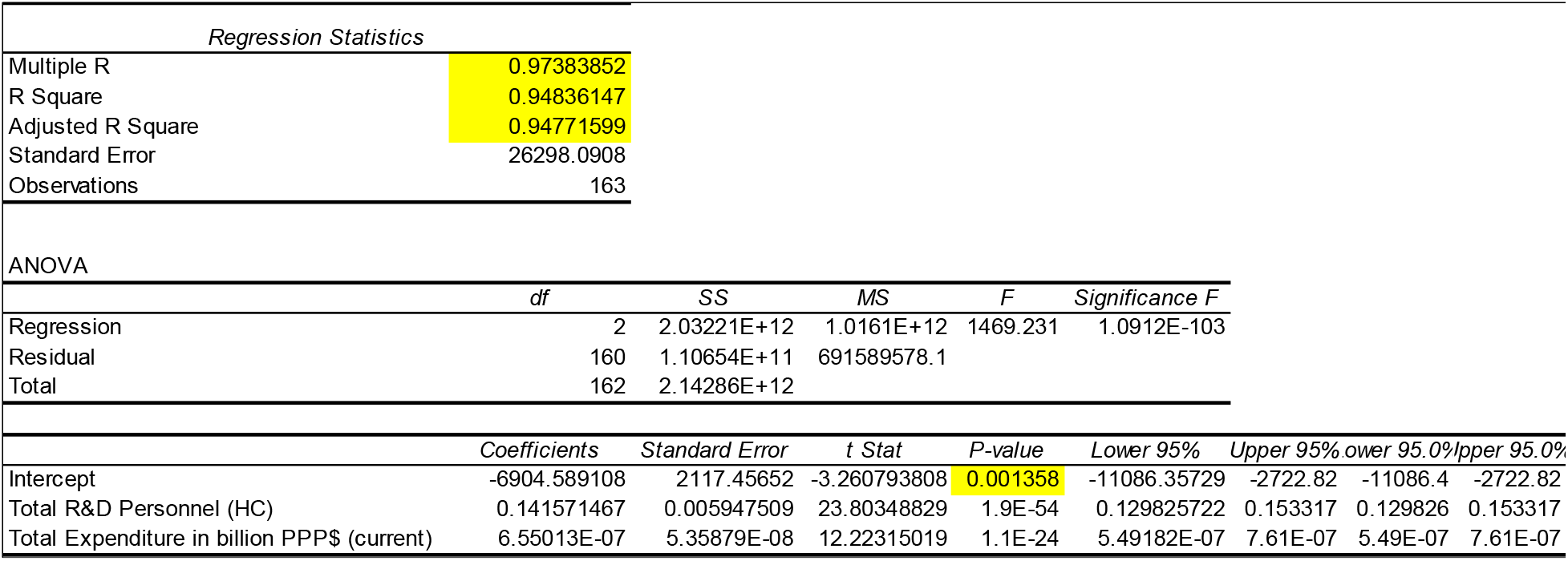
Multiple Regression analysis between No. of patent filings and Total GERD & Total R&D personnel

| Regression Statistics |  |  |  |  |  |  |  |  |
| --- | --- | --- | --- | --- | --- | --- | --- | --- |
| Multiple R | 0.97383852 |  |  |  |  |  |  |  |
| R Square | 0.94836147 |  |  |  |  |  |  |  |
| Adjusted R Square | 0.94771599 |  |  |  |  |  |  |  |
| Standard Error | 26298.0908 |  |  |  |  |  |  |  |
| Observations | 163 |  |  |  |  |  |  |  |
| ANOVA |  |  |  |  |  |  |  |  |
|  | df | SS | MS | F | Significance F |  |  |  |
| Regression | 2 | 2.03221E+12 | 1.0161E+12 | 1469.231 | 1.0912E-103 |  |  |  |
| Residual | 160 | 1.10654E+11 | 691589578.1 |  |  |  |  |  |
| Total | 162 | 2.14286E+12 |  |  |  |  |  |  |
|  | Coefficients | Standard Error | t Stat | P-value | Lower 95% | Upper 95% | Lower 95.0% | Upper 95.0% |
| Intercept | -6904.589108 | 2117.45652 | -3.260793808 | 0.001358 | -11086.35729 | -2722.82 | -11086.4 | -2722.82 |
| Total R&D Personnel (HC) | 0.141571467 | 0.005947509 | 23.80348829 | 1.9E-54 | 0.129825722 | 0.153317 | 0.129826 | 0.153317 |
| Total Expenditure in billion PPP\$ (current) | 6.55013E-07 | 5.35879E-08 | 12.22315019 | 1.1E-24 | 5.49182E-07 | 7.61E-07 | 5.49E-07 | 7.61E-07 |

**Figure 3.**
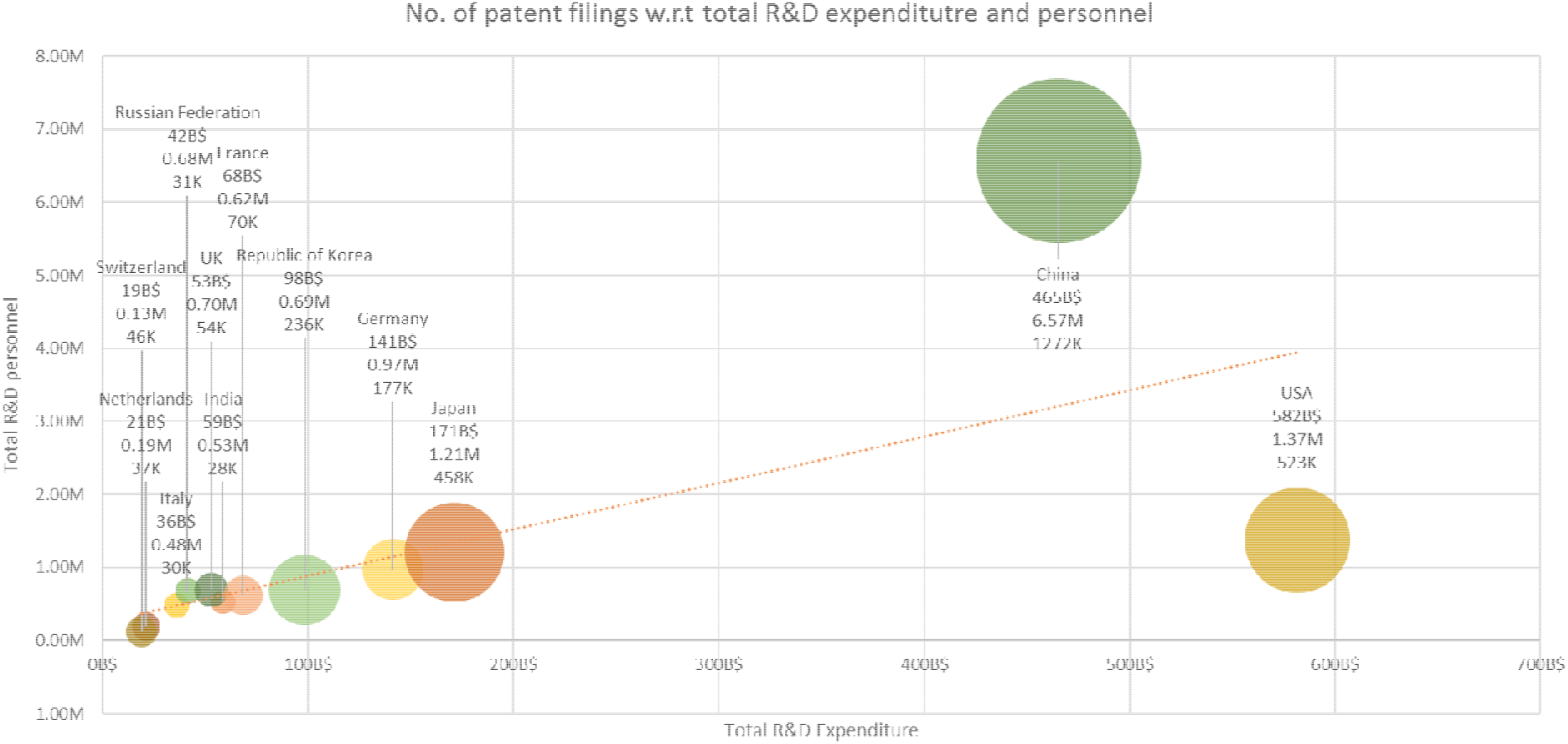
No. of patent filings with regard to total R&D expenditure and R&D personnel

However, R&D personnel includes Researchers, Technicians and supporting staff (15). We may tend to think that technicians and other supporting staff are not true researchers and exclude from the indicator but there seems to exist no or little correlation between the number of patent filings and researcher/technicians/supporting staff alone. This signifies the importance of mutual interaction and cooperation among the different stakeholders. Technicians play a major role in applying the concepts of researchers in the real world. In the top countries with the highest patent filings, technical help is providing by the industry with various levels of collaborations. The higher the researchers and technicians collaborate, the higher the patent filings from the country. Another way to say the higher the University-Industry collaboration, the higher the innovation flourishes in a country.

We can infer from the graph (Figure 4) that the countries like China, the USA, Japan, the Republic of Korea, Germany, France, Switzerland, Netherlands etc. have more patent filings than that of their scientific publications due to strong industrial collaboration. These countries have tech support in terms of skills, hardware and software, whereas those countries like Russian Federation, Italy, India, UK etc. doesn’t have strong technological support from the industry they may have fewer patent filings than that of their research publications. Though India stands 3rd position in terms of scientific research publications, most of them are confined to higher education institutions and not able to translate into real-world applications. Hence a strong collaboration with the right industry partners is the need of the moment for India to bring innovation and value addition to their products.

**Figure 4.**
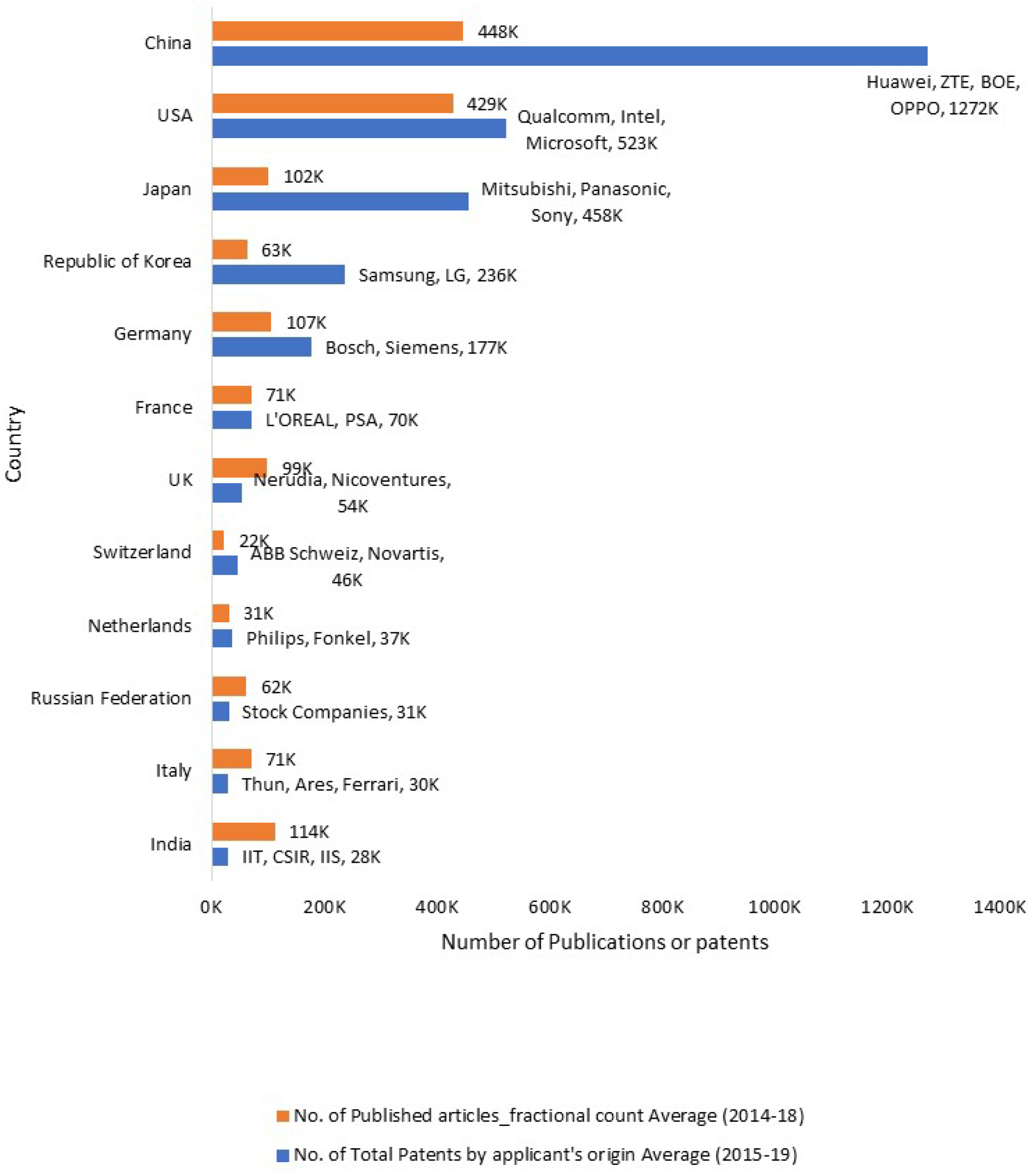
TOP 12 countries with regard to research publications and patents filings

**Figure 5.**
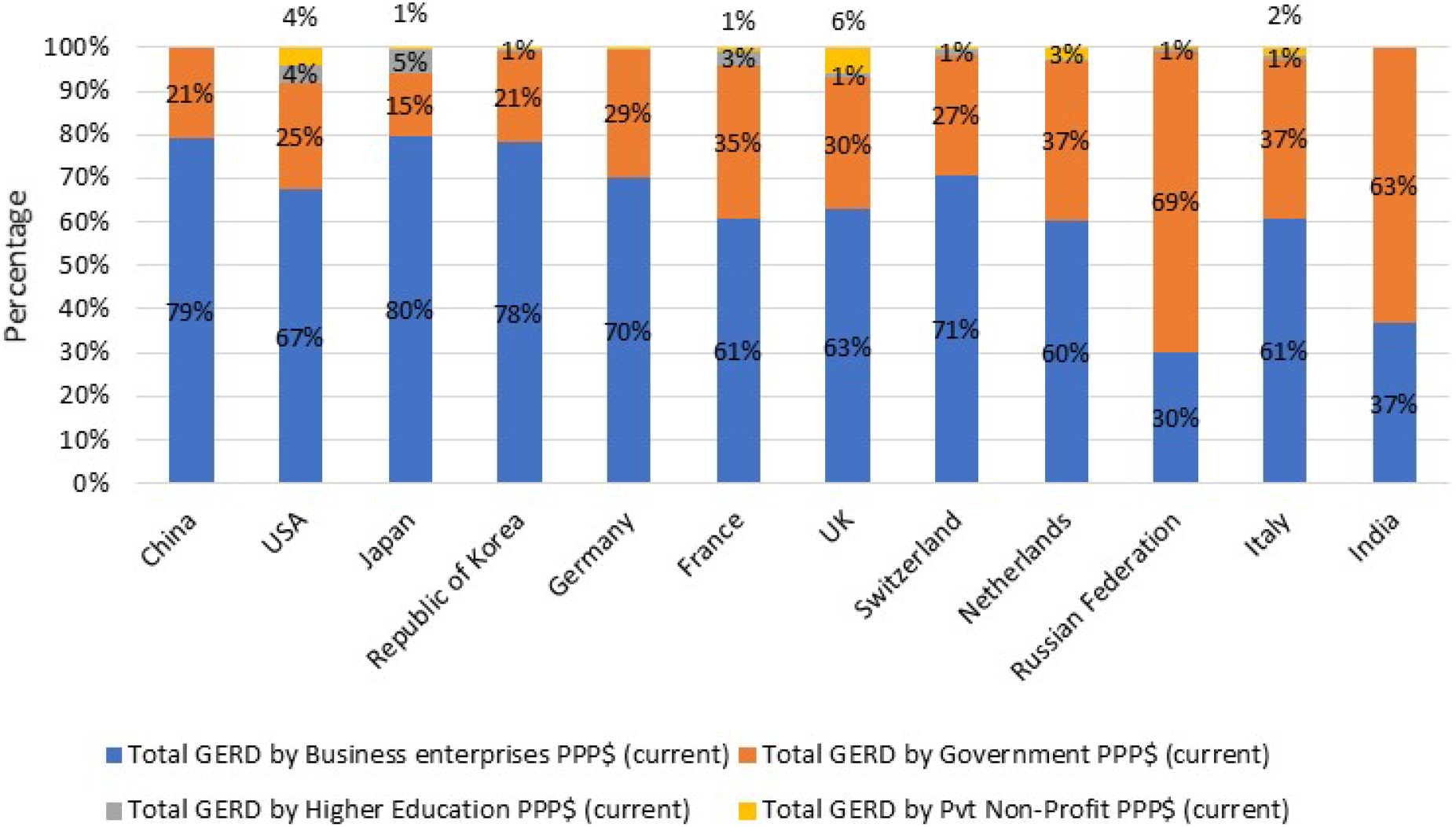
Percentage of GERD by different sectors

It can be inferred from the graph (Figure 5) that the leading countries have significant support in terms of funds from business enterprises. In contrast to the leading countries, India’s major source of funds still comes from the government and less from the business enterprises. One of the major sector where business enterprises invested in the world is pharmaceuticals (Figure 6). Though India is the largest exporter of generic drugs, R&D is being carried by only a few companies. In India, the automobile sector is the dominant one where R&D expenditure is observed but it is still far less than the other leading countries has been spending.

**Figure 6.**
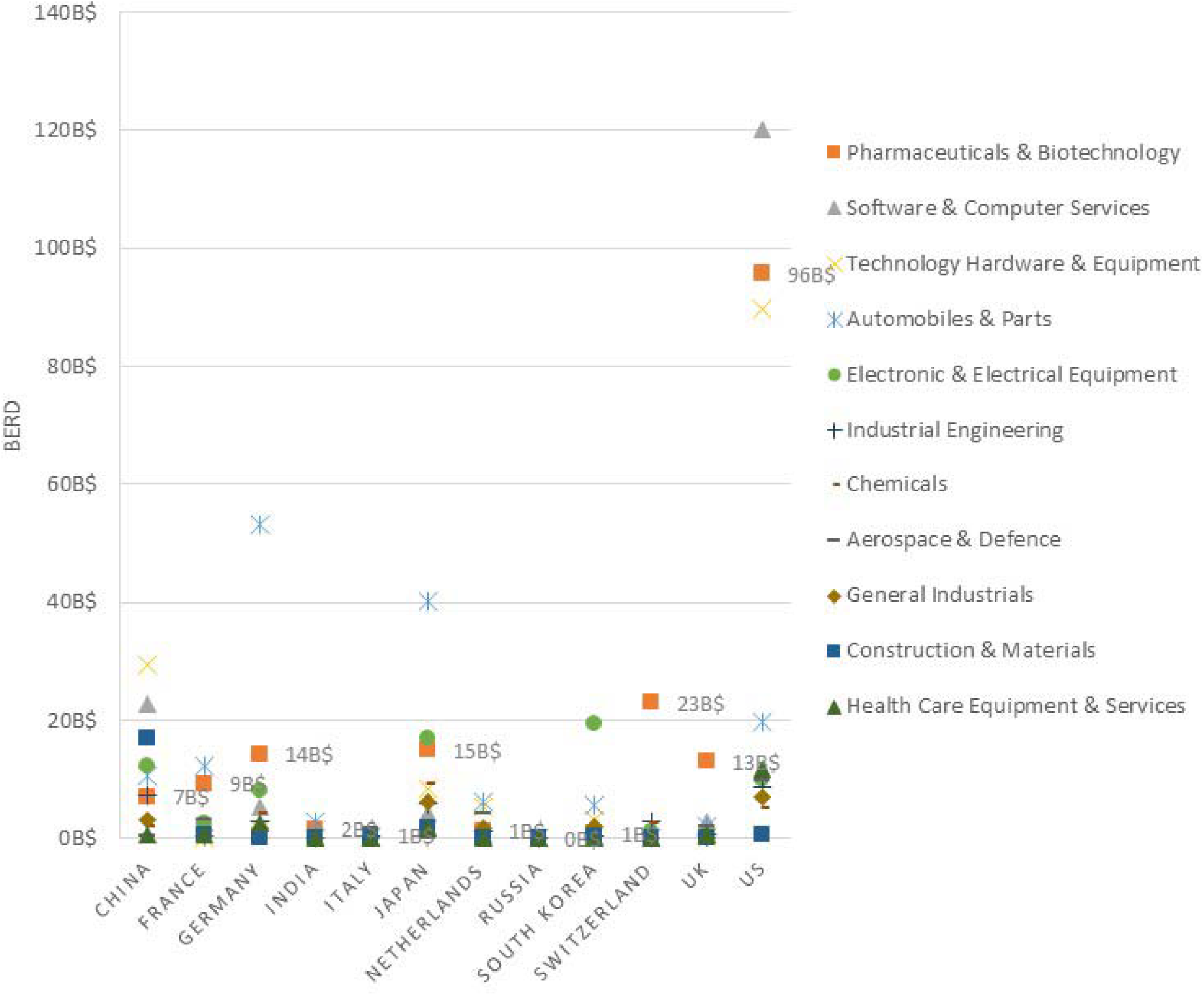
sector-wise business enterprises expenditure on R&D (BERD)

## Discussion

China had outperformed the USA and other developed nations in terms of research publications and patent filings followed by Japan and the Republic of Korea (Figure 5). The Republic of Korea has achieved unprecedented economic growth and development throughout the last four decades. (16)

Economic reforms and other related reforms in China and the Republic of Korea have changed the overall environment in which universities, the market, and the government interact (9). Over the last two decades, the concerned governments have not only initiated some major policies to reform their innovation system, but they have also set up many programs to complement these policies (9). The Republic of Korea had established PRIs (Public Research Institutions) in the 1970s and strengthened industrial R&D capabilities in the 1980s and finally established the national innovation system in the 1990s, initially they emphasized preparing a legal framework to establish innovation actors then initiated several policies for active collaboration among those innovation actors (9).

India is still in its early stages in publications and inventions compared to other leading countries. Out of 40,000 higher education institutions in India, only 1% engage in research. (The Prime Minister’s Science, Technology and Innovation Advisory Council (8). to encourage the R&D collaboration, the potential of the research institutions and universities should be strengthened by investing in R&D. It should also bring the industry into the play and collaborate with the higher education institutions to improve the skills of the human resource to enhance innovations in their respective areas (Figure 6). In 2020, India is under a way to establish NRF (National Research Foundation) to mentor research in institutions and facilitate the collaboration of researchers, government and industries around the country which is a long way to go with its limited resources (17) and without proper legal framework and policies to facilitate collaboration between industry and academia

However, if India focuses completely on collaborating with industry, it will leave less room for independence to the government and higher education sector in R&D as Industry or business enterprises have always profit motive and may focus on only those sectors which are profitable and can generate money. Industry or business enterprises may ignore other sectors like agriculture, social sciences etc. which is also reflecting in the same with countries like the Republic of Korea, Netherlands, and Russian federation. In the Republic of Korea, regardless of the source of funds and performance, engineering & Tech has occupied with predominant share and when it comes to the business enterprises it has the highest share (18).

In the case of the Netherlands, only the public sector has represented more or less equal share among all the fields of R&D but the private sector like business enterprises concentrated more on engineering & tech. In the case of Russia, the case is even worse. Business enterprises concentrated more than 90% on engineering and tech. Only the government and high education sector has shown a better share in other fields of R&D (18).

Recent research conducted in Denmark addressed the role of university-industry interaction in regional industrial development and argued in favour of a far-reaching approach in which research collaborations and graduate human capital are to be regarded as two interdependent channels. This research showed that university-industry knowledge transfer is often not immediate and, therefore, can benefit from deliberate action by both private and public actors to overcome these hurdles (19).

One also needs to consider the impact of PhD graduates and scientists’ mobility outside the academia and into regional ecosystems, as this may be regarded as a shift from knowledge transfer as a process of formal externalization and transfer of explicit and codified knowledge into more tacit and informal transfer through socialization (19).

Furthermore, any future analysis of the impact of research and development activities and knowledge transference must take into consideration not only the economically driven return of investment (ROI) but the social return of investment (SROI) (20). A discussion paper issued by the European regional office of the WHO highlights that, just like every activity, the health care sector also has an impact on the economic, environmental and social dimensions, regardless of its activities being carried out in the public, private or non-profit-making sectors and further argues that, in the spirit of the United Nations Sustainable Development Goals (SDGs), investment in healthcare must consider “their whole range of impact and directed towards sustainable and equitable solutions, which ensure health for all, leaving no-one behind” (20). Our data show that this also applies to the R&D in healthcare.

Building on the “theory of change” concept, we must define long-term goals while mapping backwards to identify necessary preconditions, thus critically designing, implementing, and evaluating initiatives and programmes intended to support changes in the original contexts by using an outcomes-based approach. (21).

In a nutshell, major players like academia and industry should collaborate effectively and efficiently to create high value-added products, and the government should frame policies to make the collaboration smooth and easy.

## List of Abbreviations

BRICS: Brazil, Russia, India, China & South Africa
FORD: Field of R&D
FTE: Full-Time Equivalent
GDP: Gross Domestic Product
GERD: Gross Domestic Expenditure on R&D
GNI: Gross Net Income
HC: HeadCount
HIC: High Income Country
LIC: Low Income Country
LMIC: Lower Middle Income Country
OECD: Organisation for Economic Cooperation and Development
NRF: National Research Foundation
PPP: Purchasing Power Parity
PRI: Public Research Institution
R&D: Research & Development
RDPPMI: R&D personnel per million inhabitants
UIS: UNESCO Institution of Statistics
UK: United Kingdom
UMIC: Upper Middle Income Country
UNESCO: United Nations Educational, Scientific and Cultural Organization.
USA: United States of America
WIPO: World Intellectual Property Organization

